# Role of sortase-assembled Ebp pili in *Enterococcus faecalis* adhesion to iron oxides and its impact on extracellular electron transfer

**DOI:** 10.1101/601666

**Authors:** Foo Kiong Ho, Ling Ning Lam, Artur Matysik, Thomas Dean Watts, Jun Jie Wong, Kelvin Kian Long Chong, Pei Yi Choo, Joe Tolar, Pui Man Low, Zhi Sheng Chua, Jason J. Paxman, Begoña Heras, Enrico Marsili, Caroline M. Ajo-Franklin, Kimberly A. Kline

## Abstract

*Enterococcus faecalis* sortase-assembled endocarditis and biofilm-associated pili (Ebp) are a virulence factor implicated in enterococcal biofilm-associated infections and gastrointestinal colonization. We previously showed that *E. faecalis* biofilm metabolism is influenced by extracellular electron transfer (EET) under iron-rich conditions, raising the question of whether Ebp pili also play a role in EET. Here, we report a novel role of Ebp pili in *E. faecalis* adhesion to the iron oxides magnetite, goethite and hematite, where the EbpA tip adhesin contributes to this interaction. Adhesion by Ebp pili is conditionally important for EET to iron oxides, as pilus mutants are attenuated in EET under non-static growth conditions. In alignment with the established role of EET in redox homeostasis, we find that EET to ferricyanide supports *E. faecalis* anaerobic growth on glycerol. Further, in an antibiotic-treated mouse gastrointestinal colonization model, we show that *E. faecalis* mutants deficient in EET poorly colonize the intestinal niche. Taken together, our findings suggest that Ebp pili can influence *E. faecalis* metabolic fitness by promoting EET to iron oxides, raising new questions of how Ebp pili shape *E. faecalis* interactions with environmental ecosystems. Furthermore, the important role of EET in *E. faecalis* colonization of the dysbiotic gastrointestinal environment highlights the need for further inquiry into how EET contributes to *E. faecalis* microbial pathogenesis.

**Importance:** In this study, we explored the interplay between extracellular electron transfer (EET) and an *Enterococcus faecalis* biofilm factor, the endocarditis and biofilm-associated pili (Ebp). We demonstrate that Ebp pili have a novel role in adhesion to iron oxides, which consequently promotes EET to iron oxides under non-static conditions. Along with our findings that *E. faecalis* EET can be coupled to anaerobic cell growth, our results point to a potential ecological role of Ebp pili in natural environments, outside of its established function in adhesion to host ligands. We provide the first evidence of the contribution of EET to *E. faecalis* colonization of the antibiotic-treated murine intestinal niche, which adds to the limited experimental evidence linking EET and microbial pathogenesis, as well as highlights the need for further studies of EET in bacterial pathogens.

## Observation

Extracellular electron transfer (EET) is a process in which microorganisms reduce extracellular substrates, such as iron, and is frequently connected to anaerobic energy metabolism (1, 2). While EET has been extensively reviewed in the Gram-negative model species *Geobacter sulfurreducens* and *Shewanella oneidensis* (3-5), the discovery of a flavin-based EET genetic locus in Gram-positive bacteria, including pathogens and members of the gut microbiota, raises the question of how microbial EET can influence human health or disease (6, 7). The gut commensal and opportunistic pathogen *Enterococcus faecalis* also performs EET (6, 8-11), but the physiological significance of EET in *E. faecalis* is not well-characterized. As our previous observations showed a relationship between *E. faecalis* biofilm metabolism and EET (8), this study focuses on investigating whether endocarditis and biofilm-associated pili (Ebp), an essential factor in *E. faecalis* biofilm formation, play a role in mediating EET. Given that EET has a prominent impact on the energy metabolism of another lactic acid bacterium *Lactiplantibacillus plantarum* (12, 13), we also sought to explore the importance of EET to *E. faecalis* using both *in vitro* and animal models.

Preliminary microscopy observations of *E. faecalis* grown in culture media autoclaved with ferric chloride showed some cells tightly surrounded by a dense material (**Fig. S1A**). This material might have been insoluble iron precipitates, inadvertently produced by autoclaving, as it was reminiscent of iron deposits previously observed in the extracellular matrix of *E. faecalis* biofilms grown under similar conditions (8). Immunofluorescence staining showed co-localization of the dense material with Ebp pili in the parent strain, while the dense material did not associate with the pilus-null Δ*ebpABC* mutant (**Fig. S1A**). As Ebp pili are associated with adhesion to fibrinogen, collagen and platelets (14, 15), as well as attachment to abiotic surfaces such as polystyrene (16), we hypothesized that there could be a similar interaction between Ebp pili and insoluble iron forms. Hence, we assessed the interaction between Ebp pili and iron by co-incubating *E. faecalis* with the iron oxide magnetite, enabling the convenient separation of iron-adherent and iron-non-adherent bacteria using a magnet (17) (**Fig. 1B**).

**Figure 1.**
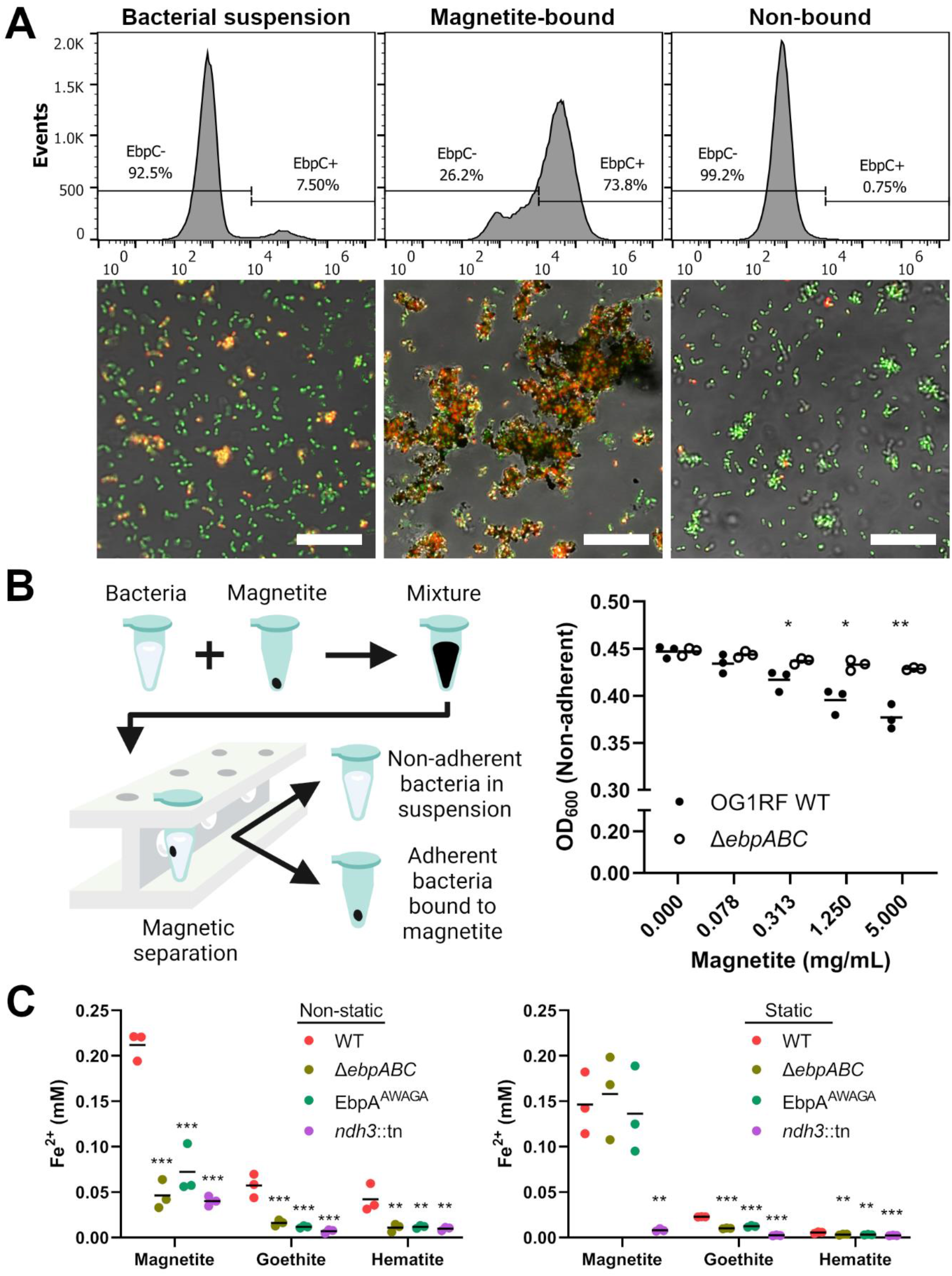
Ebp pilus expression is important for *E. faecalis* adhesion and EET to iron oxides. **(A)** Flow cytometry and immunofluorescence microscopy images of *E. faecalis* cells after magnetic separation using magnetite. Bacterial suspensions of *E. faecalis* OG1RF (left panels) were mixed with magnetite and then separated into magnetite-bound (middle panels) and non-bound fractions (right panels) using a magnetic rack. For flow cytometric analysis, the magnetite-bound bacterial fraction was released from magnetite by resuspension in 1×PBS containing 100 mM EDTA prior to staining. *E. faecalis* cells were stained with SYTO 9 DNA stain (green) and labelled for EbpC (red). Histograms show representative data from three independent experiments. Scale bars represent 20 µm. **(B)** Schematic diagram (left panel) illustrating how magnetic separation was used to isolate adherent and non-adherent fractions of bacteria. Culture turbidity (right panel) of non-adherent bacterial subpopulations after assaying with increasing concentrations of magnetite. Data points reflect optical density values at 600 nm (OD_600_) of the non-adherent fraction, after a magnet was used to remove magnetite and magnetite-bound bacteria.Horizontal lines represent the mean measurement of three independent experiments and statistical comparisons represent differences between WT and Δ*ebpABC* at each magnetite concentration. \**P* < 0.05, \*\**P* < 0.01 by unpaired Student’s *t*-test. **(C)** Iron oxide reduction by Ebp pilus mutants as assessed by the ferrozine assay under non-static (left panel) or static (right panel) growth conditions. Horizontal lines represent the mean measurement of three independent experiments. Statistical comparisons represent differences between WT and the corresponding pilus mutant. \*\**P* < 0.01, \*\*\**P* < 0.001 by one-way ANOVA with Tukey’s multiple comparisons.

Heterogenous expression of Ebp pili in *E. faecalis* OG1RF produces subpopulations of piliated and non-piliated cells (14-16, 18). Through immunofluorescence microscopy and flow cytometry, we observed that the pilus-expressing cell population selectively adheres to magnetite (**Fig. 1A**; **Fig. S1B**). The Δ*ebpABC* mutant showed significantly less adherence to magnetite compared to the parental wild-type strain (**Fig. 1B**; **Fig. S2**). Using the EbpA^AWAGA^ mutant, which has a mutated metal ion-dependent adhesion site (MIDAS) motif to disrupt EbpA function without perturbing pilus biogenesis (19), we found that the tip adhesin EbpA contributes to magnetite binding (**Fig. S2C**). Magnetite binding was also attenuated in the single deletion mutants of EbpA and major fiber pilin EbpC, while no significant differences were observed for the deletion mutant in cell wall anchor pilin EbpB (**Fig. S2C**). Findings from single deletion mutants overall correlate with previously published biofilm formation phenotypes due to defective pilus biogenesis (20). We were also interested in investigating whether the adhesion phenotype was similar for goethite and hematite, both of which are amongst the most common iron oxides found in soils (21). To test for adhesion to other iron oxides, we separated non-adherent bacteria from mineral-bound bacteria using differential density centrifugation with a sucrose solution and similarly observed adhesion to goethite and hematite in a pilus-dependent manner (**Fig. S2D**).

To determine if the adhesin function of Ebp pili influences EET to iron oxides, bacteria were grown in the presence of iron oxides and the amount of ferrous iron produced and released into the supernatant was quantified by the ferrozine assay. As the control, we used an EET-deficient mutant containing an insertional inactivation of the *ndh3* gene. *E. faecalis* Ndh3 is a NADH dehydrogenase that had previously been characterized to be a component of the electron transport chain involved in EET, where it oxidizes NADH and transfers electrons to demethylmenaquinone (6, 9). Ndh3 is distinct from Ndh2, which is likely the NADH dehydrogenase specific for aerobic respiration (9). Significant differences in iron reduction were observed between parental and pilus mutant strains when bacterial growth was conducted under non-static conditions, while these differences were absent or less distinct under static growth conditions (**Fig. 1C**). Due to the heme auxotrophy of *E. faecalis* and the absence of heme in the culture media, observations made here were likely not influenced by the effects of aerobic respiration. Collectively, our findings indicate that Ebp pili play an accessory role in enhancing EET through the aggregation of *E. faecalis* cells with iron oxides. Though we did not differentiate between direct and indirect electron transfer, the results suggest that close contact between bacteria and iron oxides is important for efficient electron transfer. It is also tempting to speculate that the binding of Ebp pili to iron oxides is an evolutionary trait that augments bacterial EET in environments, such as soils and sediments, where iron oxides are commonly present. Ferrous iron produced by EET might additionally modulate pilus function by competition with or displacement of the native divalent cation in the MIDAS motif of EbpA (**Fig. S3**).

Previous work in *Listeria monocytogenes* demonstrated that the electron transport chain involved in EET can route electrons not only to extracellular iron, but also through extracellular reductases such as fumarate reductase (22). As anaerobic glycerol dissimilation in *E. faecalis* is dependent on fumarate reductase (23), we predicted that EET could provide an alternative route to external electron acceptors in place of fumarate. To test if the membrane-impermeable ferricyanide could act as the electron acceptor for EET during glycerol dissimilation, we grew *E. faecalis* macrocolonies anaerobically on an agar base supplemented with ferricyanide. To limit excess fermentation of amino acids, growth medium was excluded from the agar base and instead, the bacterial inoculum was resuspended in growth medium prior to spotting on agar plates. We find that the presence of either fumarate or ferricyanide promotes *E. faecalis* growth on glycerol, whereas growth was deficient in the absence of an electron acceptor or when the EET-deficient *ndh3*::tn mutant was used (**Fig. 2A**). Our findings confirmed that *E. faecalis* EET can be coupled to anaerobic cell growth, which is consistent with metabolic and growth phenotypes implicated in EET for *L. monocytogenes* and *L. plantarum* (6, 13). It would be particularly interesting to make further comparisons against *L. plantarum*, as it is a lactic acid bacterium similar to *E. faecalis* and has been shown to combine features of fermentation and respiration in a hybrid form of metabolism (13). As EET also promoted *L. monocytogenes* colonization of the mouse gastrointestinal tract (6), we investigated whether *E. faecalis* EET plays a similar role. As anticipated, lower numbers of *ndh3*::tn mutant were recovered in an antibiotic-treated mouse gastrointestinal model of colonization when compared to the parent strain (**Fig. 2B**).

**Figure 2.**
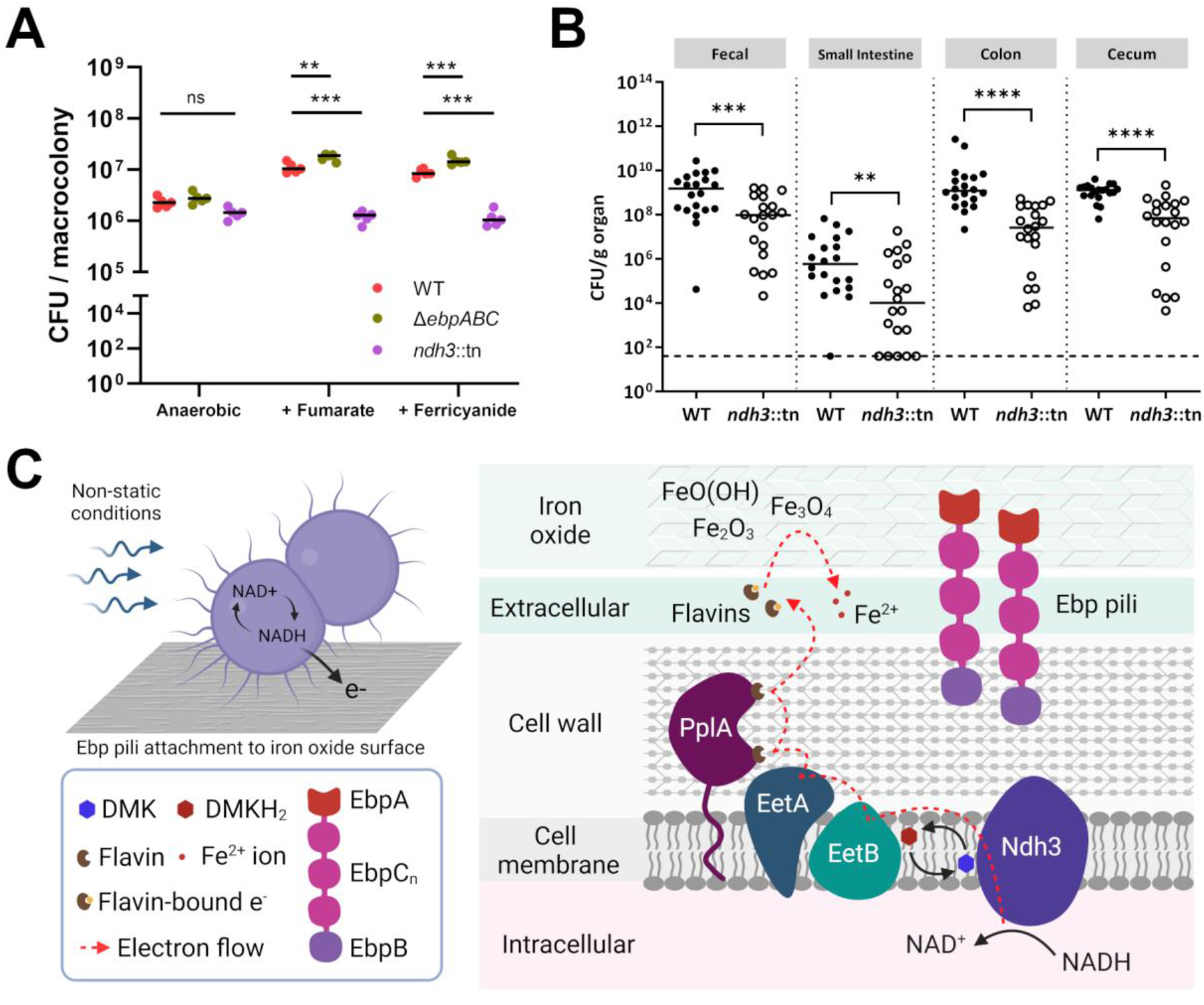
*E. faecalis* EET contributes to optimal fitness *in vitro* and *in vivo*. **(A)** Anaerobic macrocolony growth with glycerol as the carbon source in the presence of either 2.5 mM fumarate or 2.5 mM ferricyanide as the electron acceptor. After 24-hour growth in an anaerobic jar, macrocolonies were excised from the agar and CFU enumerated. Data shown are from five independent experiments and each dot represents the average of three technical replicates. Horizontal lines represent the median CFU. Statistical comparisons were assessed by one-way ANOVA with Tukey’s multiple comparisons. \*\**P* < 0.01, \*\*\**P* < 0.001, ns non-significant. **(B)** *E. faecalis* colonization of lower gastrointestinal tract using an antibiotic-treated mice model of gut colonization. Datapoints represent individual mice and the dotted line indicates the limit of detection at CFU <40. Data shown are from four independent experiments with 5 mice per group in each experiment. Horizontal lines represent the median CFU. Statistical comparisons were assessed by the Mann-Whitney test. \*\**P* < 0.01, \*\*\**P* < 0.001, \*\*\*\**P* < 0.0001, ns non-significant. **(C)** Proposed model for accessory role of Ebp pili in promoting adhesion and EET to iron oxides. Efficient EET to iron oxides under non-static conditions is mediated by Ebp pili, which has a novel role in adhesion to magnetite, goethite and hematite. Ebp pili are not essential for EET, but adhesion promotes close contact with iron oxides for more efficient EET. The tip pilin EbpA, which is responsible for adhesion to host collagen and fibrinogen, contributes to the adhesion to iron oxides but it is not known if the binding mechanism is similar for these substrates. EET contributes to the maintenance of redox homeostasis (NAD^+^/NADH balance), which is known to be important for various metabolic processes such as the anaerobic glycerol dissimilation pathway in *E. faecalis*. While lactate fermentation is the primary mode of NAD^+^ regeneration in *E. faecalis*, experiments in another lactic acid bacterium *L. plantarum* have demonstrated that EET supplements fermentative pathways to increase metabolic flux and yield. Lastly, the physiological consequence of EET to iron oxides would be highly dependent on the thermodynamic favorability of electron transfer in these environments, but a possibility is that EET serves as a strategy for *E. faecalis* to maintain competitive fitness by expanding the range of electron acceptors that it is able to utilize. Components are not drawn to scale. Figure is created using BioRender.

In summary, we describe a novel role of *E. faecalis* Ebp pili in adherence to iron oxides, which may promote EET and enhance metabolic fitness in environmental ecosystems where iron oxides are present (**Fig. 2C**). While we show that EET can be coupled to anaerobic growth of *E. faecalis*, it is possible that EET may play additional roles, such as in iron uptake or detoxification (24, 25). Iron availability is a strong driver for colonization of the gastrointestinal tract by pathogens (26), but it is still unclear how Ebp pili will interact with different iron forms present in the gastrointestinal tract. Nevertheless, we demonstrate that mutants deficient in EET poorly colonize the mouse gastrointestinal tract, indicating that EET contributes to *E. faecalis* outgrowth during gut dysbiosis.

## Acknowledgments

This work was supported by the National Research Foundation and Ministry of Education Singapore under its Research Centre of Excellence Programme and by the Ministry of Education Singapore under its Tier 2 programme (MOE2014-T2-2-124). Foo Kiong Ho and Artur Matysik were supported by Singapore Ministry of Health’s National Medical Research Council under its Open Fund Individual Research Grant (MOH-000645) and its Clinical Basic Research Grant (NMRC/CBRG/0086/2015), respectively. Begoña Heras and Jason Paxman were supported by the Australian Research Council (ARC) project grant (DP210100673) and the National Health and Medical Research Council (NHMRC) project grant (GNT1143638). We thank Hailyn V. Nielsen and Scott Hultgren (Washington University, St Louis, USA) for providing the pilus mutants, and Gary Dunny (University of Minnesota, USA) for providing transposon mutants. We also thank Bala Davient and Caroline Manzano for their assistance with mutant construction.

## Data Availability

All data generated or analysed during this study are included in this published article (and its supplementary information files).

