## Supplementary information for "Role of sortase-assembled Ebp pili in *Enterococcus faecalis* adhesion to iron oxides and its impact on extracellular electron transfer"

**This file includes:**

**Supplementary Materials and Methods**

**Supplementary Figures S1-S3**

**Supplementary Table 1**

**Supplementary References**

### Supplementary Materials and Methods

**Bacterial strains and growth conditions.** *E. faecalis* strains were routinely maintained on Brain Heart Infusion (BHI; Becton, Dickinson and Company, Franklin Lakes, NJ) agar plates. *E. faecalis* overnight cultures were grown in Tryptic Soy Broth (TSB; BD Bacto, USA) and cultured at 37°C under static conditions. When sterility was required, magnetite, goethite and hematite powder (all from Sigma-Aldrich, USA) were immersed in 70% ethanol, followed by three washes in sterile 1×PBS, then diluted to desired concentrations. For preliminary experiments of *E. faecalis* biofilm formation under iron-supplemented conditions, culture medium was prepared by autoclaving TSB broth supplemented with 0.175% glucose and 2 mM $FeCl_3$. Bacteria were grown as biofilms in 6-well plates for 24 h at 37°C, then disrupted with a cell scraper in 1×PBS prior to downstream analysis.

**Magnetite binding assays.** Overnight cultures of *E. faecalis* strains were centrifuged and normalized in 1×PBS to an optical density value at 600 nm ($OD_{600}$) of 0.5. 900 µL of normalized bacterial suspension were mixed with either 100 µL of magnetite suspension or with 100 µL of 1×PBS, followed by brief mixing with a vortex for association of bacteria to magnetite particles. Magnetite particles were removed from the mixture by placing samples on a magnetic rack and transferring the supernatant to a new tube. The resulting supernatant, which contained the non-adherent bacteria, was transferred to a cuvette and measured in a UV-Vis spectrophotometer at $OD_{600}$. The amount of bacterial adherence to magnetite can be estimated by the loss of $OD_{600}$ in bacterial suspensions mixed with magnetite compared to bacterial suspensions that were mixed with 1×PBS only. For CFU enumeration, magnetite particles were washed three times in 1×PBS to remove non-adherent bacteria and resuspended in a solution of 1×PBS containing 100 mM EDTA, adjusted to pH 7.4. Samples were vortexed vigorously to dissociate adherent bacteria from magnetite particles, followed by separation of magnetite particles from the supernatant using a magnet. The number of bacteria that had adhered to magnetite was estimated by CFU plating of supernatant on BHI agar plates. For crystal violet assays, magnetite particles were washed three times in 1×PBS to remove non-adherent bacteria and the adherent biomass was stained for 10 minutes with a solution of 0.1% crystal violet prepared in 1×PBS. After staining, the magnetite particles were transferred to a new tube to reduce background reading and then washed five times in 1×PBS with vortexing. The crystal violet stain was then eluted from adherent biomass using 96% ethanol. Magnetite particles were removed with a magnet and the supernatant was measured in a spectrophotometer at 590 nm.

**Flow cytometry and fluorescence microscopy.** Bacterial fractions were prepared by magnetic separation as described above and normalized to an $OD_{600}$ of 0.5. Bacterial cells were blocked in 2% (w/v) bovine serum albumin (BSA) in 1×PBS for 1 h at room temperature (RT). Bacterial cells were then incubated for 1 h at RT with guinea pig α-EbpC serum (1), washed 3 times with 1×PBS and stained for another hour at RT with goat anti-guinea pig secondary antibody Alexa Fluor 647 (Invitrogen, USA). Both α-EbpC serum and secondary antibody were diluted at a 1:500 ratio in 2% BSA/PBS. Bacterial cells were subsequently stained with Syto9 DNA stain (Invitrogen, USA), diluted at a 1:500 ratio in 1×PBS, before washing twice with 1×PBS. Flow cytometry was performed using the BD Accuri C6 Flow Cytometer (BD Biosciences, USA), where at least 10,000 events were collected for each condition. FlowJo version v10 software (BD Biosciences, USA) was used for analysis and plotting of flow cytometry data. Fluorescence imaging was performed on the Elyra PS.1 LSM780 system and the Axio Observer Z1 system (Carl Zeiss, Göttingen, Germany).

**Iron mineral reduction assays.** Ferric reductase activity on insoluble iron minerals was assessed using the ferrozine assay, which was performed as previously described with several modifications (2). Bacterial cultures were initially grown to mid-log phase in TSB broth and then normalized to $OD_{600}$ of 0.5 in sterile 1×PBS. Normalized cultures were centrifuged and resuspended in TSB broth supplemented with 0.4% glucose (TSBG), followed by a further 1:100 dilution in TSBG broth to prepare the inoculum. The assay was initiated in 1.5 mL microcentrifuge tubes by mixing 500 µL of inoculum with 500 µL TSBG supplemented with the indicated iron minerals. Final concentrations of magnetite were at 1 mM, goethite at 3 mM and hematite at 1.5 mM to ensure that an equivalent molar concentration of Fe atoms was added to each reaction. After a brief vortexing, tubes were either incubated at 37°C under static conditions or in a thermomixer set to shake at 15-minute cycles comprised of 1 minute shaking and 14 minutes rest. After 24 hours of incubation, samples were centrifuged at 8,000 g for 5 minutes and the supernatant was transferred to a new tube. 200 µL of supernatant was added to 800 µL of 2 mM ferrozine in 1×PBS for measurement of ferrous iron. After 5 min of colour development in the dark, the absorbance of samples was measured in a spectrophotometer at 562 nm with appropriate dilutions. Concentration of iron in the supernatant was calculated against a standard curve prepared using $FeSO_4$ with 10 mM L-cysteine HCl dissolved in 1×PBS.

**Estimation of adhesion by sucrose differential density centrifugation.** Procedures to separate adherent bacteria from non-adherent bacteria was adapted from another study (3). *E. faecalis* strains were grown overnight in TSB (BD Bacto, USA) broth at 37˚C under static conditions. Bacterial-mineral mixtures were prepared by combining 900 µL of normalized bacterial suspensions with 100 µL of mineral suspension and vortexed briefly. Mineral suspensions were prepared by adding an amount equivalent of 0.001 $cm^3$ of mineral to each tube. 1 mL bacterial-mineral mixture was then overlaid on top of a 70% sucrose solution in 1×PBS, followed by centrifugation in a swinging bucket rotor at 3,000 g for 5 minutes. The supernatant was then aspirated from the top to remove any non-adherent bacteria. The pellet, containing minerals and associated bacteria, was resuspended in 1×PBS and transferred to a new microcentrifuge tube. The tube was centrifuged, and the pellet was resuspended in TE (10 mM Tris-HCl, 1 mM EDTA) buffer prior to transferring into Lysing Matrix B tubes. Bead-beating was performed at 6 m/s for 40 seconds for 2 cycles to lyse adherent bacteria that was associated with minerals. Tubes were centrifuged at 16,000 g for 5 minutes to sediment the debris and the clear lysate was transferred to a new tube. The amount of released DNA in the lysates was quantified as a relative measure of the amount of bacteria using the Qubit dsDNA High-Sensitivity assay (Invitrogen, USA), according to manufacturer's instructions.

**EbpA Structural Modelling.** The EbpA amino acid sequence was submitted to the structural homology-modelling server Swiss-Model (https://swissmodel.expasy.org) (4). Template structures for structural modelling were automatically selected based upon sequence identity. The initial template search derived 723 templates that were filtered down to 50 templates, whereby 14 models were constructed and ranked based on sequence similarity and coverage. The top solution was based upon the PDB 3txa (GBS104 from *Streptococcus agalactiae*) that showed 34% sequence similarity and 38% sequence coverage. The final

EbpA model included N177-P620. The $Fe^{2+}$ was created with Sketcher (5) and placed into position using the MIDAS bound magnesium of 3txa and 2ww8 as a guide with the program COOT (6). The two conserved water molecules from both 3txa and 2ww8 were fitted into EbpA by a similar means. Final alignments, analysis and structural figures were created with PyMOL (7).

**Modified macrocolony assay for EET-dependent growth.** An agar base containing the tested electron acceptor was prepared by mixing one part of 3% agar with one part of 5 mM potassium ferricyanide (Sigma-Aldrich, USA) or 5 mM sodium fumarate dibasic (Sigma-Aldrich, USA), where all components were dissolved in 1×PBS. 20 mL of this mixture was dispensed into each Petri dish and then allowed to solidify. Bacterial suspensions at $OD_{600}$ of 0.5 were prepared as described above and were inoculated into culture medium at a 1:100 ratio. Culture medium composed of TSB without dextrose (Sigma-Aldrich, USA) supplemented with 2% glycerol as the carbon source. 5 µL of inoculated cultures were spotted onto the prepared agar base in triplicates and incubated at 37°C for 24 hours in an anaerobic jar. After incubation, macrocolonies were excised from the agar base and transferred into a microcentrifuge tube containing 1 mL of 1×PBS, followed by thorough vortexing to resuspend the bacteria. Serial dilution was performed, and the appropriate dilutions were enumerated for CFU by spread plating on BHI agar plates.

**Mouse Gastrointestinal Tract (GI) Infection Model.** Three-week-old male C57BL/6NTac mice were administered ampicillin (VWR, USA) in their drinking water (1 g/L) for 5 days as previously described (8). Mice were then given one day of recovery from antibiotic treatment prior to administration of approximately 1-5 x $10^8$ CFU/mL *E. faecalis* ($OD_{600}$ of 0.5) in the drinking water for 3 days. Before and after infection, mice were monitored for signs of disease and weight loss. All animal experiments were approved and performed in compliance with the Nanyang Technological University Institutional Animal Care and Use Committee (IACUC). At the indicated timepoints, the small intestine, colon, and cecum were harvested. Tissue samples were homogenised and serial diluted 1×PBS prior to spot-plating on BHI agar with 10 mg/L colistin, 10 mg/L nalidixic acid, 100 mg/L rifampicin, 25 mg/L fusidic acid for CFU enumeration. All antibiotics were obtained from Sigma Aldrich, USA.

### Supplementary Figures

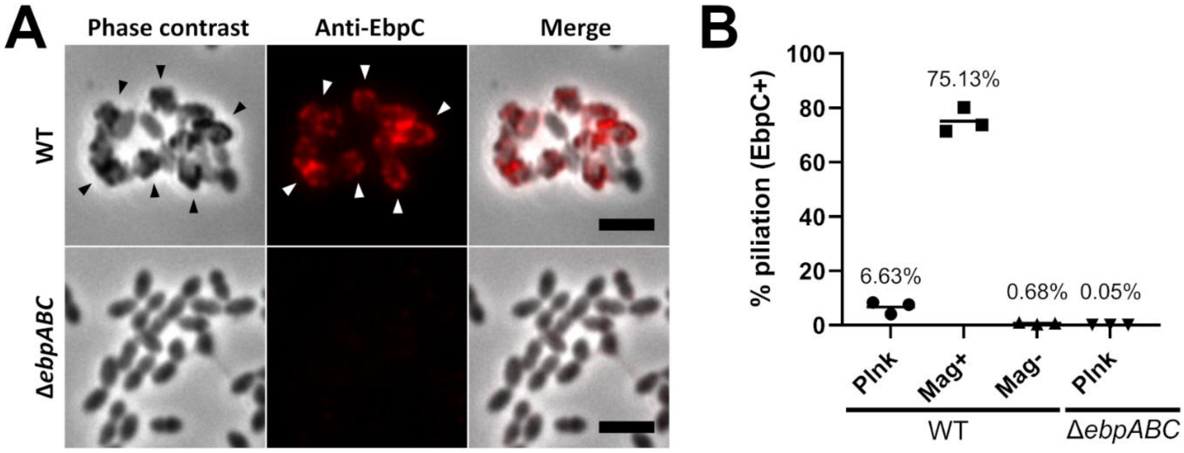


**Figure S1. Insoluble iron attaches to *E. faecalis* expressing Ebp pili. (A)** Ebp pili-expressing *E. faecalis* accumulates a dense material on its cell surface when grown in culture medium autoclaved in the presence of ferric chloride. Images show disrupted *E. faecalis* biofilm cultures labelled for EbpC (red). Representative microscopic images from three independent experiments are shown. Arrowheads show instances where a dense material observed under phase contrast co-localizes with fluorescent anti-EbpC staining. Scale bars represent 3 µm. **(B)** Corresponding plot for the flow cytometry data in **Fig. 1A**, showing the proportions of Ebp pili-expressing bacteria after magnetic separation using magnetite. Horizontal lines and percentage values represent the mean measurement of three independent experiments. X-axis represents the following conditions: Plnk, unfractionated planktonic cultures; Mag+, magnetite-bound fraction; Mag-, non-bound fraction.

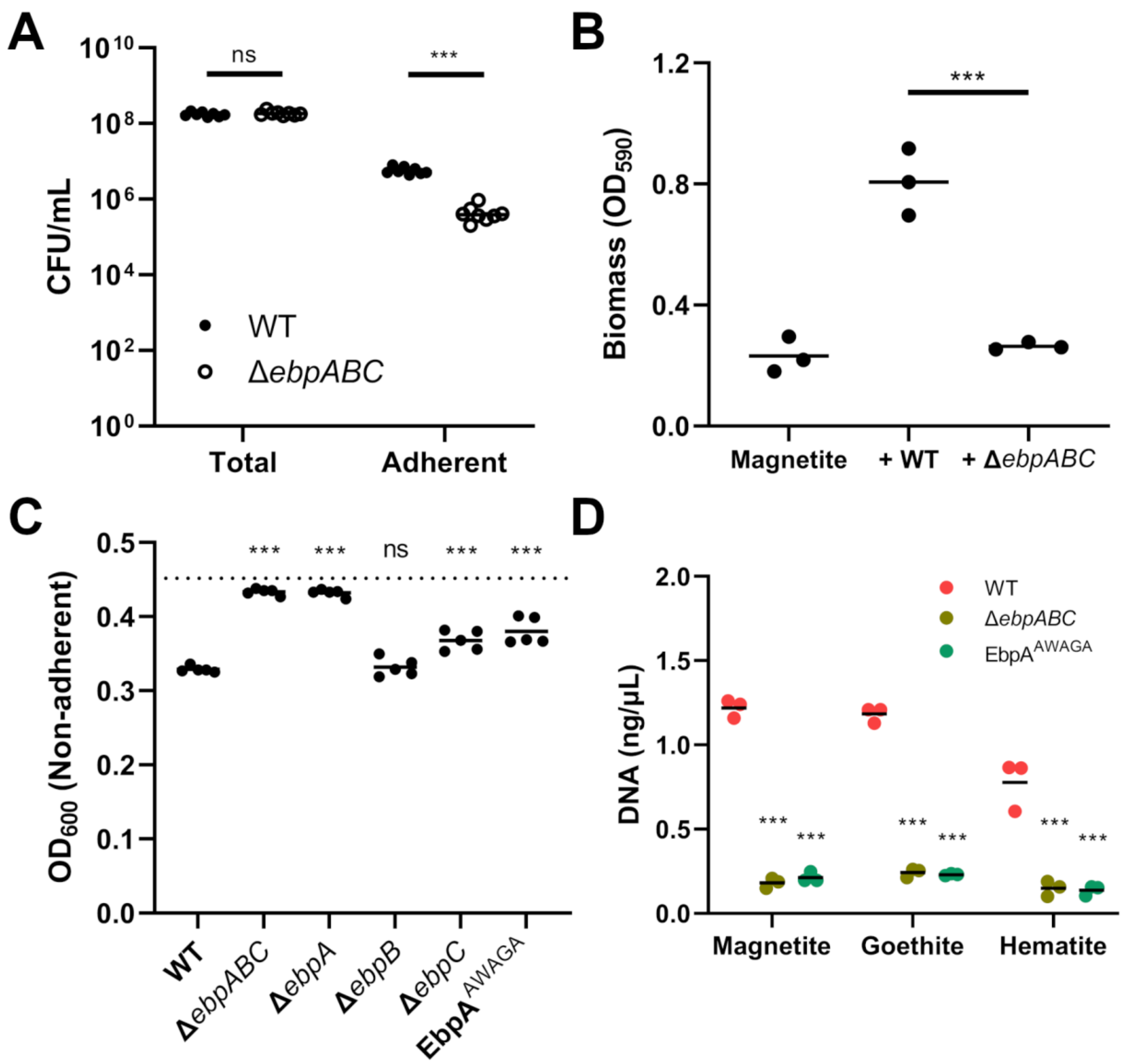


**Figure S2. The tip adhesin EbpA contributes to adhesion to iron oxides. (A)** CFU enumeration of the magnetite-bound bacterial fraction after dissociation using 1×PBS containing 100 mM EDTA. Horizontal lines represent median CFU. ****P* < 0.001, ns non-significant, by Mann-Whitney test. **(B)** Estimation of biomass adherent to magnetite determined by crystal violet assay. Data points for the first column represent crystal violet staining of magnetite without addition of bacterial strains. Horizontal lines represent the mean measurement. ****P* < 0.001, ns non-significant by one-way ANOVA with Tukey's multiple comparisons. **(C)** Culture turbidity of non-adherent subpopulations of indicated Ebp pilus mutants assayed with 5 mg/mL magnetite. Data points above reflect optical density values at 600 nm ($OD_{600}$) after a magnet was used to remove magnetite and magnetite-bound bacteria. Horizontal dotted line represents the mean culture turbidity for bacterial suspensions with no addition of magnetite. Statistical comparisons represent differences between WT and the corresponding pilus mutant. ***P* < 0.01, ****P* < 0.001, ns non-significant, by one-way ANOVA with Tukey's multiple comparisons. **(D)** Adherent and non-adherent bacteria were separated by sucrose differential density centrifugation. Data points above reflect supernatant DNA concentrations, as a relative measure of adherent bacteria, after the adherent bacterial fractions were lysed by bead-beating. Horizontal lines represent the mean measurement. Data shown are from at least three independent experiments. ****P* < 0.001 by one-way ANOVA with Tukey's multiple comparisons.

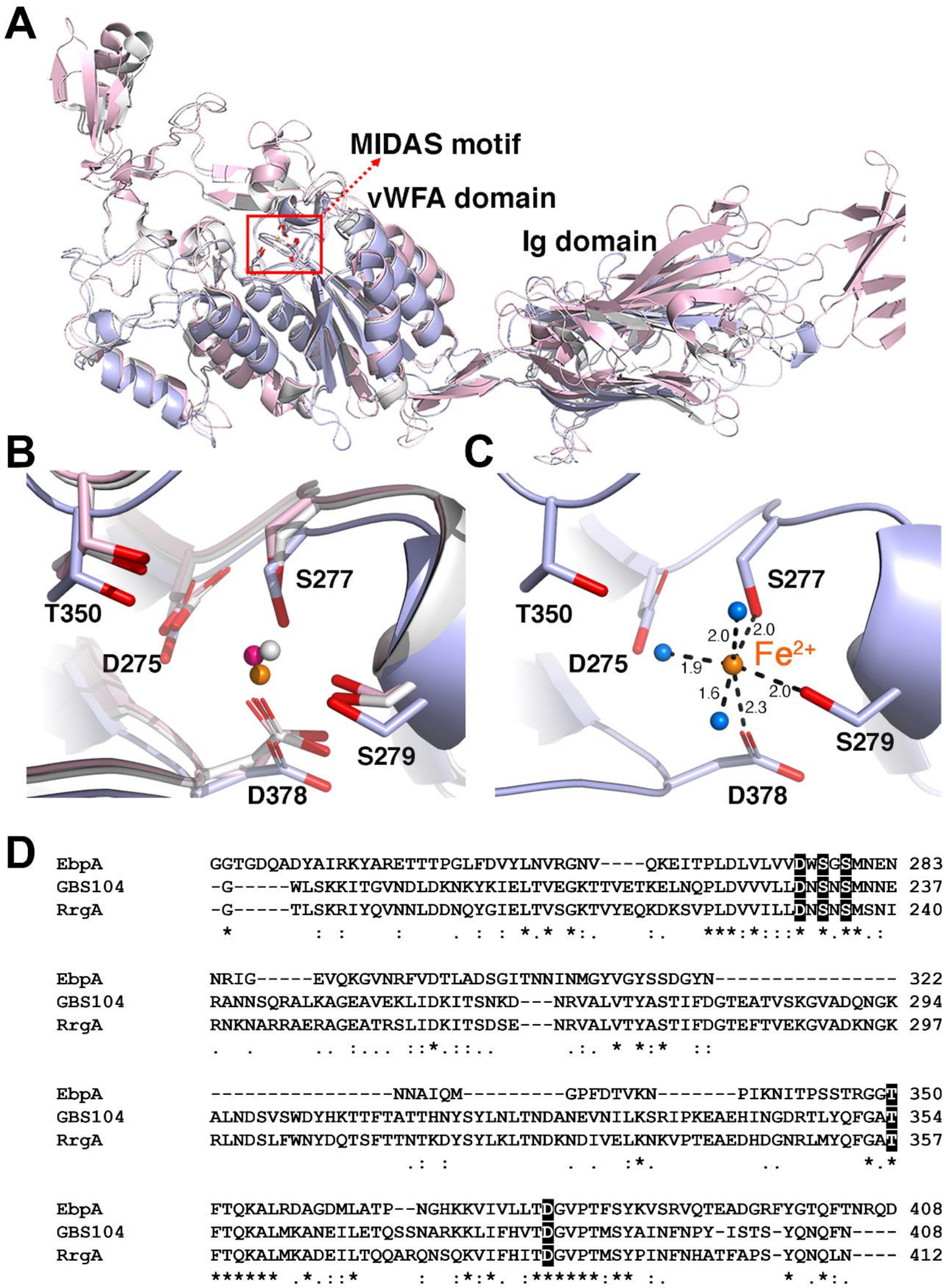


**Figure S3. The EbpA MIDAS motif may be a potential binding site for ferrous iron. (A)** Superimposition of EbpA structural model (N177-P620) (blue) with that of homologs *S. agalactiae* GBS104 (grey) and *S. pneumoniae* RrgA (pink). In agreement with other pilin adhesins, EbpA tip folds into a vWFA domain comprising of a 5 stranded β-sheet surrounded by 7 α-helices. The MIDAS motif is located at the N-terminus of the β-sheet. Only a partial model was obtained for the 2 extended arms of this domain. The EbpA tip is then followed by an IgG fold domain. **(B)** Closeup view of the superimposition of the MIDAS motif from EbpA (blue), GBS104 (grey) and RrgA (pink) showing conservation of D275, S277, S279, T350 and D378 residues (EbpA numbering). Bound $Mg^{2+}$ ions (grey/magenta) in GBS104 and RrgA and modelled $Fe^{2+}$(orange) in EbpA are shown as spheres. **(C)** The $Fe^{2+}$ is coordinated by the EbpA MIDAS motif by ~2Å direct interactions with S277, S279, D378 and a water-mediated interaction with D275 along with interactions with two additional water molecules (water molecules shown as blue spheres). Upon a closed-to-open conformational change, bonding could shift from D378 to T350. **(D)** Clustal alignment of the EbpA sequence with homologs *S. agalactiae* GBS104 (Q8E0S5) and *S. pneumoniae* RrgA (A0A0H2UNT6), showing conservation of the EbpA MIDAS motif and associated metal-binding residues. Residues from G228 to D408 (EbpA numbering) are displayed.

**Supplementary Table 1. Strains used in this study.**

| Strain | Description | Reference |
|---|---|---|
| OG1RF | Laboratory strain; $Rif^R$, $Fus^R$ | (9) |
| Δ*ebpABC* | *ebpABC* chromosomal deletion mutant; $Rif^R$, $Fus^R$ | (10,11) |
| Δ*ebpA* | *ebpA* chromosomal deletion mutant; $Rif^R$, $Fus^R$ | (10,11) |
| Δ*ebpB* | *ebpA* chromosomal deletion mutant; $Rif^R$, $Fus^R$ | (10,11) |
| Δ*ebpC* | *ebpC* chromosomal deletion mutant; $Rif^R$, $Fus^R$ | (10,11) |
| $EbpA^{AWAGA}$ | EbpA allelic replacement with $EbpA^{AWAGA}$; $Rif^R$, $Fus^R$ (coding for $Ala^{315}$-Trp-$Ala^{317}$-Gly-$Ala^{319}$) | (10,11) |
| *ndh3*::tn | Transposon insertion mutant of *ndh3*; $Rif^R$, $Fus^R$, $Cm^R$ | (12,13) |
